# Analysis of selection signatures reveals important insights into the adaptability of high-altitude Indian sheep breed *Changthangi*

**DOI:** 10.1101/2020.12.05.412916

**Authors:** Sheikh Firdous Ahmad, Arnav Mehrotra, Sona Charles, Nazir Ahmad Ganai

## Abstract

*Changthangi* is a high-altitude sheep breed of India that is adapted to cold and hypoxic climate of Himalayas. In the present study, we analysed population structure of *Changthangi* and contrasted it with selected Indian and European commercial sheep breeds to detect genomic regions under positive selection. The studied domesticated sheep breeds included *Changthangi, Indian Garole, Deccani, Tibetan, Rambouillet* and *Australian Merino*. While the PCA results revealed *Changthangi* clustered with *Tibetan* sheep; TREEMIX and ADMIXTURE results also detected the introgression of lowland Indian sheep inheritance in *Changthangi*. Cross-population comparisons of *Changthangi* using XP-EHH showed multiple functional regions present on OAR 7, 15 and 16, to be under selection in *Changthangi* sheep. These regions are related to adaptation to climatic and hypoxic stressors, nervous system and mitochondrial functioning. The genes present in these regions are suitable candidates for future studies on the genetic mechanisms underlying high-altitude adaptation.

## Introduction

Sheep is one of the earliest domesticated farm animal species and current sheep population has evolved over thousands of years of domestication (Pedrosa et al., 2005). Natural and artificial selection for various traits (production, reproduction and other functional traits) has resulted in phenotypic adaptations which may be behavioural, physiological and/ or psychological. Domestication stamps detectable signatures on the phenome and genome of modern livestock breeds in comparison to wild counterparts of a particular species or a population (Gutiérrez-Gil et al., 2017). Selection results in increased gene frequency of favoured alleles and produces hitchhiking effect on frequency of neutral alleles which are present at linked loci (Smith and Haigh, 2008). Heterogeneity in environment (including climate and precipitation) also influences the spatial distribution and other aspects of phenotypic and genetic variation in livestock and other organisms (Lv et al., 2014). Elucidation of these signatures at molecular, and more recently at genomic levels, provides an opportunity in animal breeding to understand genetic basis of different adaptive phenotypes and reciprocally how genetic variations affect phenotypes. Improved accessibility to whole-genome data and advances in bioinformatics analysis has revolutionized our understanding of major biological pathways under selection.

India is one of the mega-biodiversity hotspots with a large livestock inventory. It is home to more than 74 million sheep and 148 million goats reared in different agro-climatic conditions. The rearing conditions vary from high mountainous terrains of Himalayas to marshy Sunderbans (Eastern India) to plains of northern and central India. Domestication in varied agro-climatic zones and selection for different objectives has given rise to the development of 44 sheep breeds (http://www.nbagr.res.in/regsheep.html) that are registered with Indian nodal agency *i.e*., National Bureau of Animal Genetic Resources (NBAGR). The important breeds maintained at varied conditions include *Changthangi, Garole, Deccani, Tibetan, Chokla, Kendrapada, Marwari, Muzaffarngri, Poonchi, Karnah, Nellore* and others. Few exotic breeds have traditionally been used in breeding policy of India for sheep which include *Rambouillet, Australian Merino, Poll Dorset* and *Corriedale*. *Changthangi* sheep breed is reared by Changluk community in Ladakh region of India and is famous for its adaptation to high altitude conditions (feed shortage, low oxygen) with extremely cold climate during winters (−20 to −40 °C) and adversities of snow (Ganai et al., 2011). The breeding tract consists of rugged mountainous terrains and is situated at an altitude of 14-16 thousand feet above mean sea level. The animals are reared by nomads at highlands with high altitude areas and rarely brought to low hills for grazing purposes.

In sheep, several studies have tried to explore the selection signatures associated with different phenotypes ranging from morphological traits to production, adaptation and disease resistance (Kim et al., 2016; Lv et al., 2014; McRae et al., 2014; Moioli et al., 2013; Moradi et al., 2012). We hypothesized that adaptation to the harsh high-altitude conditions has left signatures of selection on the *Changthangi* genome. Also, no previous study has investigated the regions under selection in any high-altitude Indian sheep breed. Therefore, our objective in this study was to study the population structure of the *Changthangi* breed and to contrast the breed with selected Indian and European commercial sheep breeds to detect the genomic regions under selection through cross-population extended haplotype homozygosity (XP-EHH) method.

## Material and methods

### Data information

The present study utilized the genome-wide Illumina OvineSNP50v1 data on 292 animals from seven different sheep breeds. The domesticated sheep breeds included in the study were *Changthangi* (n=29), *Garole* (n=26), *Deccani* (n=24), *Tibetan* (n=37), *Rambouillet* (n=102) and *Australian Merino* (n=50). *European Mouflon* (n=24) was used as an out-group for studying the stratification and phylogenetic lineage of breeds under study as it is considered as progenitor of major global sheep population of the globe (Hiendleder et al., 2002). The genotypic data were retrieved from the WIDDE database (Kijas et al., 2012; Sempéré et al., 2015). The SNP coordinates corresponded to the Oar_v3.1 assembly.

The breeds chosen for comparison are reared under varied agro-climatic conditions and selection programmes are applied for different traits. The main purpose of their domestication varies from fine wool to dual purpose (both meat and wool). Among the wool-purpose breeds, *Australian Merino* produce fine wool (average fibre diameter less than 20 microns) while the yield from *Changthangi* sheep is mainly carpet wool (average fibre diameter of 41 microns). *Changthangi* and *Tibetan* are dual purpose breeds and both of them are reared under high altitude conditions, though the altitudes and climatic conditions differ significantly. On the other hand, *Garole* and *Deccani* sheep belong to mutton-purpose breeds that are reared in plain areas and selected exclusively for growth and meat production. *Rambouillet* is a dual purpose exotic breed with its origin in New Zealand. According to agro-ecological classification of Indian sheep breeds, the breeds under study belonged to three different classes i.e., north-temperate (*Changthangi*); southpeninsular (*Deccani*); and eastern (*Garole* and *Tibetan*).

### Quality control and LD pruning

From the merged dataset of all the breeds, unmapped and non-autosomal SNPs were filtered out. Subsequently, Plink v1.9 (Purcell et al., 2007) was used to apply quality control parameters as minor allele frequency (--maf 0.01), Hardy-Weinberg equilibrium (--hwe 0.000001), and SNP missingness (--geno 0.1), leaving 46,819 SNPs for downstream analysis. Furthermore, the markers in 50 Kb non-overlapping windows with linkage disequilibrium greater than 0.1 were identified and removed (–indep-pairwise 50 5 0.1) for Admixture and TREEMIX analysis.

### Population stratification

We examined the population stratification of seven breeds under study using three different approaches *i.e*., principal component analysis (PCA), model-based genomic clustering, and TREEMIX analysis. PCA approach helps to visualize the relationship among individuals of same and different breeds based on genotype matrix. For PCA with all 292 animals, the genomic relationship matrix (GRM) was constructed using GCTA v 1.93 (Yang et al., 2011) and the top 20 principal components (PCs) of the GRM were calculated. Plotting of various PCs was undertaken using R programming environment.

In model-based analysis, ancestry coefficients are used to infer population stratification. ADMIXTURE v1.3 program (Alexander and Lange, 2011) was used for model based genomic clustering and genomic ancestry analysis with *K* values of 2-7. Analysis was done for five iterations per *K* value using random seeds and crossvalidation estimates were recorded for each run for the different *K* values. The results were plotted using PONG (Behr et al., 2016). TREEMIX is another model-based approach that provides an opportunity to study splits and admixtures in population structure and substructure. TREEMIX software (Pickrell and Pritchard, 2012) was run on the LD pruned data, using *European Mouflon* as the root of the tree in the analysis to study population splits and mixtures.

### Identification of selection sweeps

We attempted to detect the selection signatures in *Changthangi* sheep breed which could have resulted from its adaptation to high altitude, cold climate and acute feed shortage at times, especially during winters. *XP-EHH* method was employed to detect selection sweeps for the different breed-combinations as *Changthangi* against each breed individually; *Changthangi* against all Indian breeds combined (*Tibetan*, *Deccani* and Indian *Garole*) and against the European commercial breeds combined (*Australian Merino* and *Rambouillet*); thus making a total of seven combinations. First, the haplotypes were phased using Beagle 5.1 (Browning et al., 2018) for all the breeds. XP-EHH scores were then calculated for *Changthangi’s* comparison with each of the 7 breed groups using ‘rehh’ R-package (Gautier and Vitalis, 2012). P values were derived by transforming the XP-EHH scores into - log[Φ(XPEHH)] in which Φ(XPEHH) is the cumulative Gaussian distribution function. Pxpehh >= 4 was considered as significant. Subsequently, candidate regions were identified using a 100 kb scanning window with 50 kb overlap. Genes in the significant regions were obtained from Ensembl 101 genes database. The human orthologs of the genes under positive selection in *Changthangi* were used for functional profiling using g:Profiler server (Reimand et al., 2016). The protein-protein network among these genes was constructed using STRINGv11 (Szklarczyk et al., 2019).

## Results and discussion

### Population stratification

#### Principal component analysis

To study the population stratification, three different approaches were employed. PCA is aimed to reduce the dimensionality of the genomic relationship matrix so that underlying individuals (and breeds) are separated along different principal components (PCs). The first three principal components were able to explain much of the variation (17.5% of total variation among breeds under study) and generated near-exclusive clusters based on *indicine-exotic* inheritance or in breed-wise fashion (Fig. 1A and 1B). Principal component 1 (PC1) differentiated the breeds under study into three main non-overlapping clusters *i.e*., Cluster#1 (the outgroup; *European Mouflon*); Cluster#2 (European breeds; *Rambouillet* and *Australian Merino*) and cluster#3 (*Indicine* breeds; *Changthangi*, *Deccani*, *Tibetan* and *Garole*). No inter-cluster overlapping was observed. However, overlapping was observed among *indicine* sheep breeds within the same cluster. PC3 stratified different Indian sheep breeds under study with *Garole* being farthest from others. In PCA plots, *Changthangi* clustered closest to *Tibetan* sheep. The separation of clusters corresponded to the geographical origin of the sheep breeds under study.

**Fig. 1:**
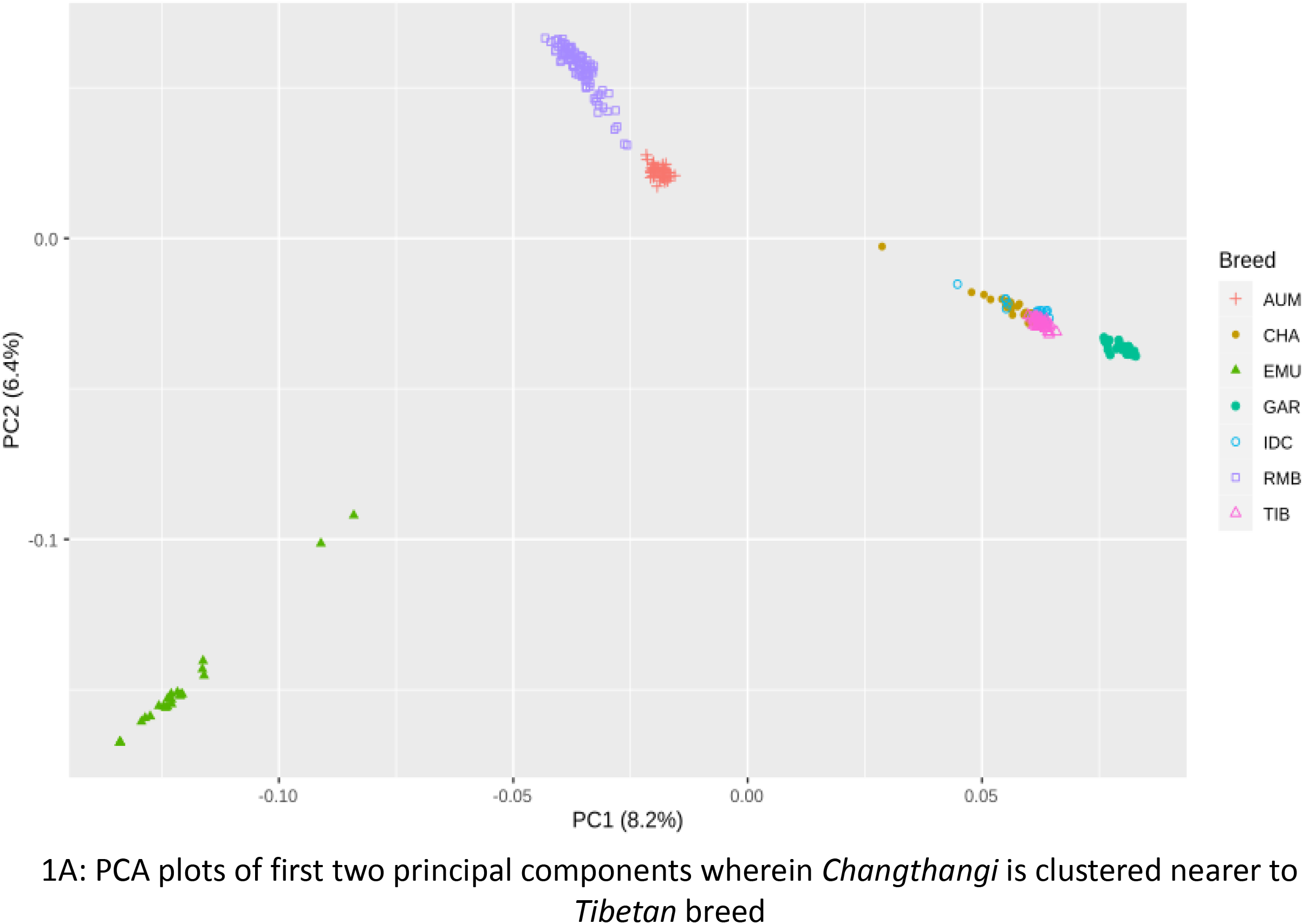
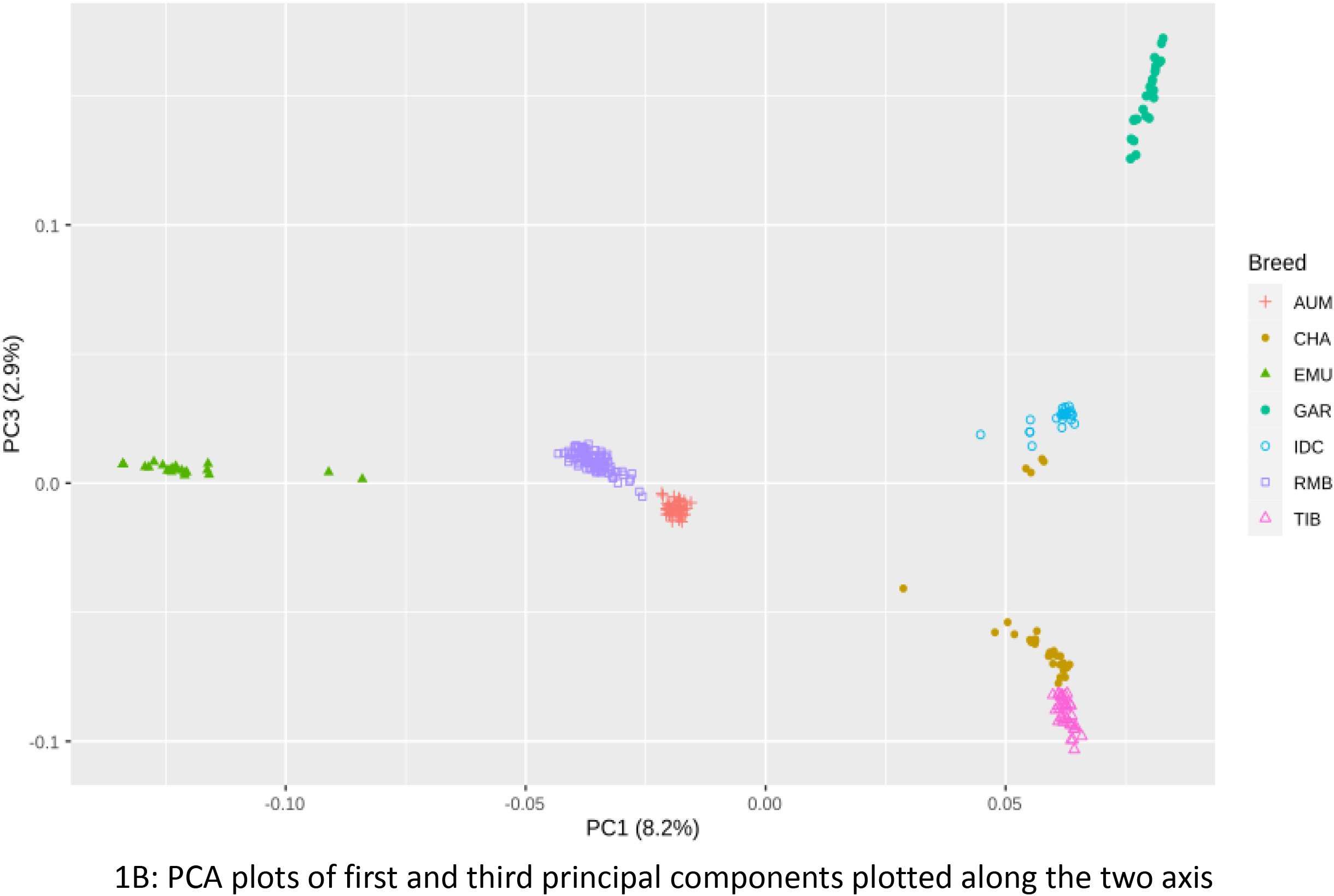
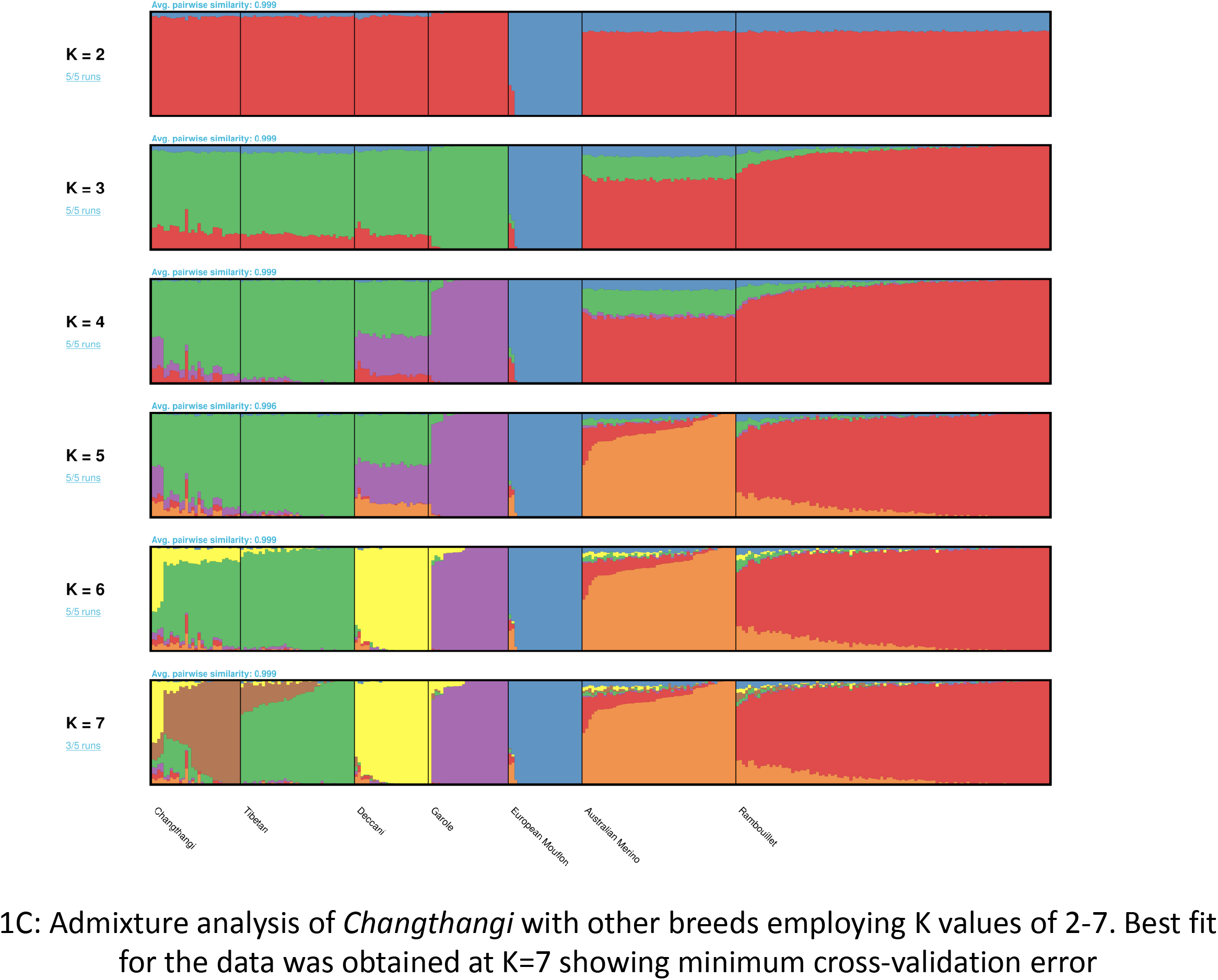
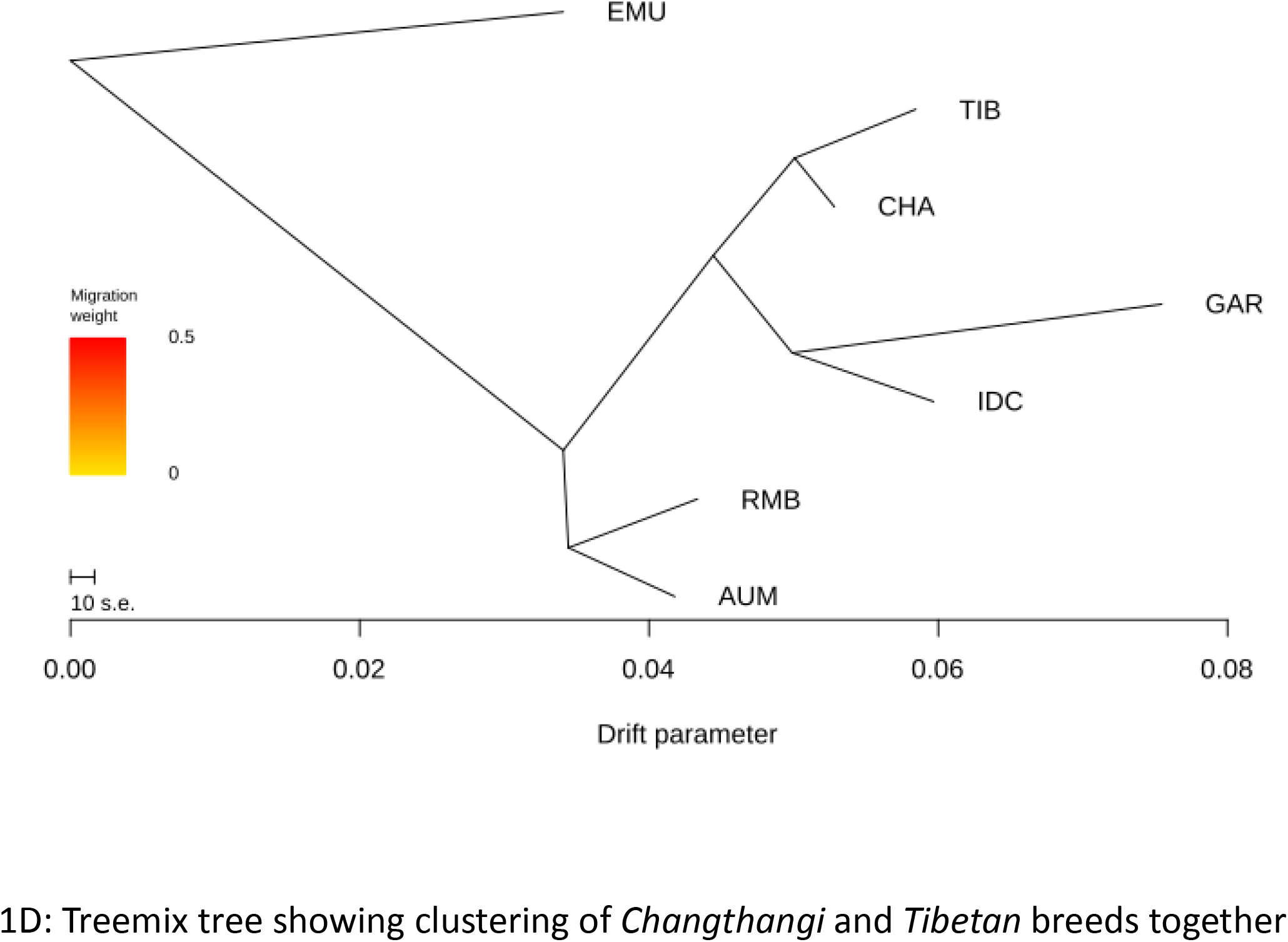
Population structure of *Changthangi* breed on comparison with other breeds.

#### Admixture analysis

To further explore the population structure among individuals of different breeds under study, model-based hierarchical clustering analysis was undertaken for *K*-values of 2-7 (*K*-being the user-defined number of biological ancestral populations). Cross-validation (CV) estimates revealed *K*-value of 7 to be best fit for the breeds under study; it scored minimum CV error among all *K* values. The barplot with CV error estimates plotted against each K-value is presented in Fig. 1C. Analysis with *K*=2 separated *European Mouflon* from the domesticated breeds. Separation between the *indicine* and European breeds was clear at K=3. At K=6, *Changthangi* shared high amount (69.1%) of inheritance with Tibetan sheep. It has been proposed that the present day *Changthangi* and *Tibetan* breeds have originated from a common group (Gorkhali et al., 2016). However, the presence of 19.8% shared inheritance with a lowland Indian breed-*Deccani* remained unexplained. At the *K*-value of 7, each breed was assigned its own cluster. The complex mixture of *Tibetan* and *Deccani* inheritance, with traces of European ancestry was visible in *Changthangi*.

#### Treemix analysis

The phylogram with no migrations, generated *via* TREEMIX analysis, explained 99.5% of the observed covariance between the breeds (Fig. 1D). *Rambouillet* and *Australian Merino* formed a separate clade, as did *Garole* and *Deccani*. The high altitude breeds *i.e., Changthangi* and *Tibetan* were clustered together in phylogram. Addition of a single migration edge showed introgression of *Deccani* inheritance in *Changthangi* and explained 99.8% covariance. This was concordant with the presence of *Deccani* inheriance in Admixture results, and the clustering of few *Changthangi* animals near *Deccani* in the PCA plot. It may be possible that *Deccani* or another lowland Indian sheep with shared ancestry with *Deccani* was introduced in the high-altitude areas of Ladakh in the past. However, we could not find any records documenting such a cross-breeding attempt. Nevertheless, our findings demonstrate the need to thoroughly investigate the breed composition and the breeding history of *Changthangi* population.

#### Selection signatures

XP-EHH is considered as a powerful procedure for detection of signatures of ongoing selection (Sabeti et al., 2002) and is based on probability of homozygosity in given extended haplotypes from different populations which carry a specific core haplotype structure (Wei et al., 2016). Under *XP-EHH* procedure, EHH scores are estimated from extended haplotypes containing same core haplotype and comparison is made among populations. Positive and negative XP-EHH score are indicative of selection sweeps in population under consideration and reference group, respectively (Wei et al., 2016).

Selection sweeps were analysed for a total of 7 combinations involving *Changthangi* and different Indian and European breeds. A summary of the SNPs and regions identified in each combination is given in the Table 1. We found 35 genes under positive selection in *Changthangi* across all comparisons (Table 1). The protein-protein interaction network of the genes is given in supplementary file S1.

The Manhattan plots depicting the genome-wide distribution of significant selection sweeps in *Changthangi* breed on comparison with other breeds are presented in Fig. 2A-G. Results revealed a 720 Kb segment on Chromosome 15 (50.98 Mb – 51.70 Mb) bearing SNPs under positive selection in *Changthangi* (pXP-EHH > 4) in comparisons with Indian as well as European breeds. Three different comparisons highlighted this region; *Australian Merino* – with the top SNP s13677.1 (XP-EHH = 5.22); *Tibetan* – with the top SNP s53691.1 (XP-EHH = 4.05), and the combined Indian breed group with the top SNP s71974.1 (XP-EHH = 5.02). A peak at this region could also be observed in the combined comparison with European breeds (*Merino* and *Rambouillet*) but the top SNP s63809.1 fell just short (XP-EHH = 3.89) of the set threshold. A search in Ensembl database showed 17 genes were present in this region which included *P2RY6, ARHGEF17, RELT, FAM168A, U6, PLEKHB1, RAB6A, MRPL48, COA4, PAAF1, DNAJB13, UCP2, UCP3, C2CD3* and *PPME1*.

**Fig. 2:**
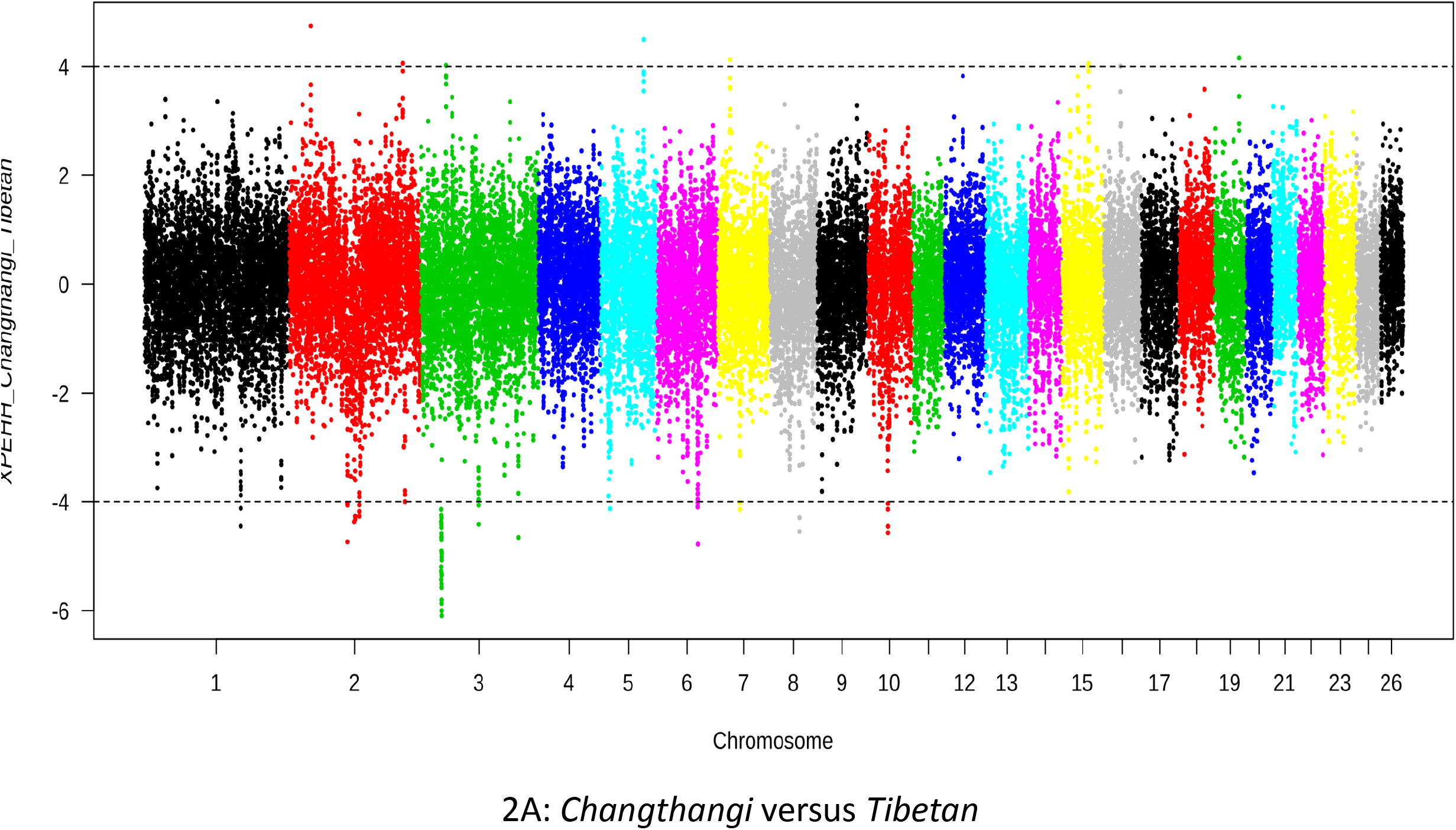
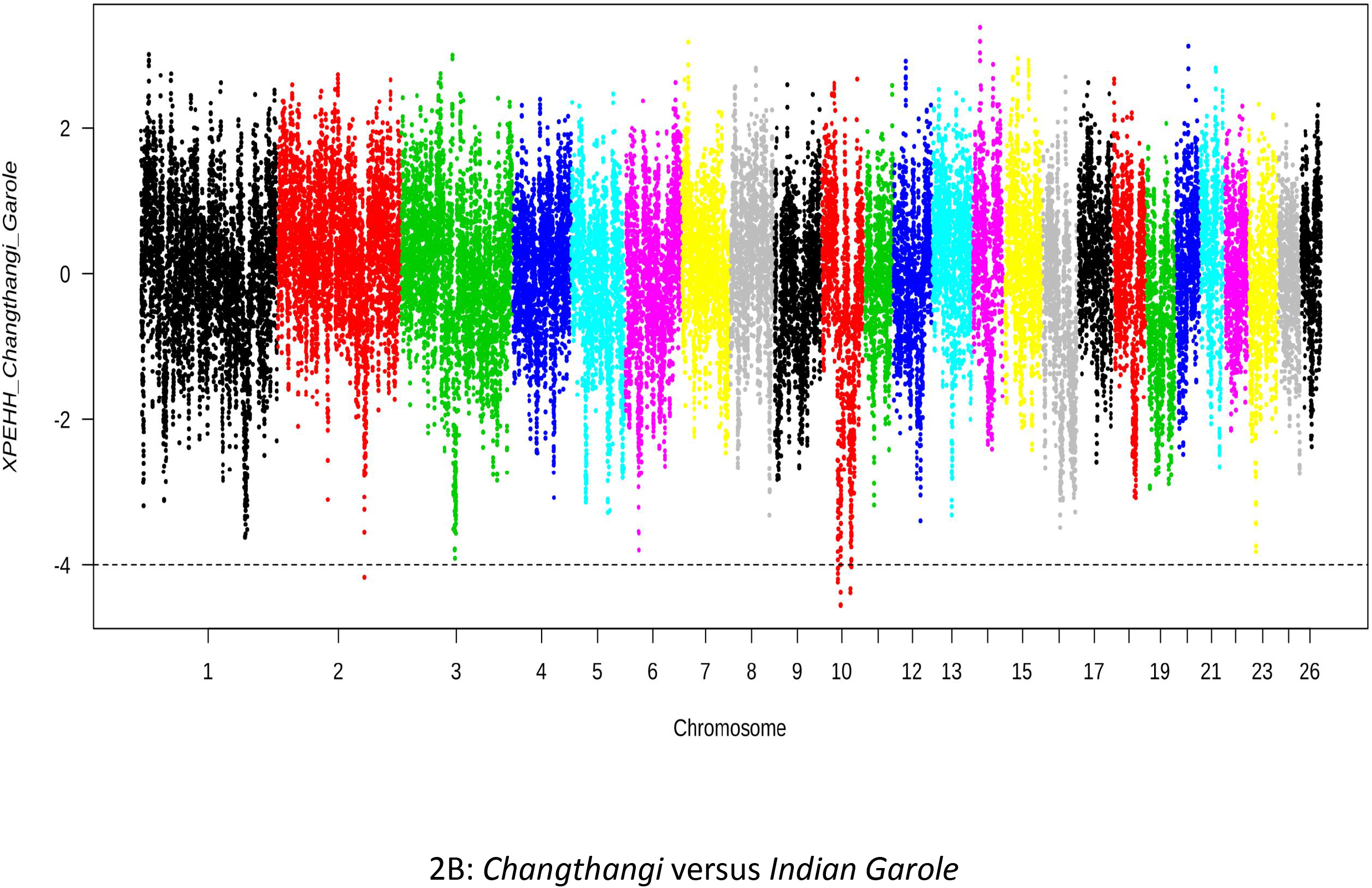
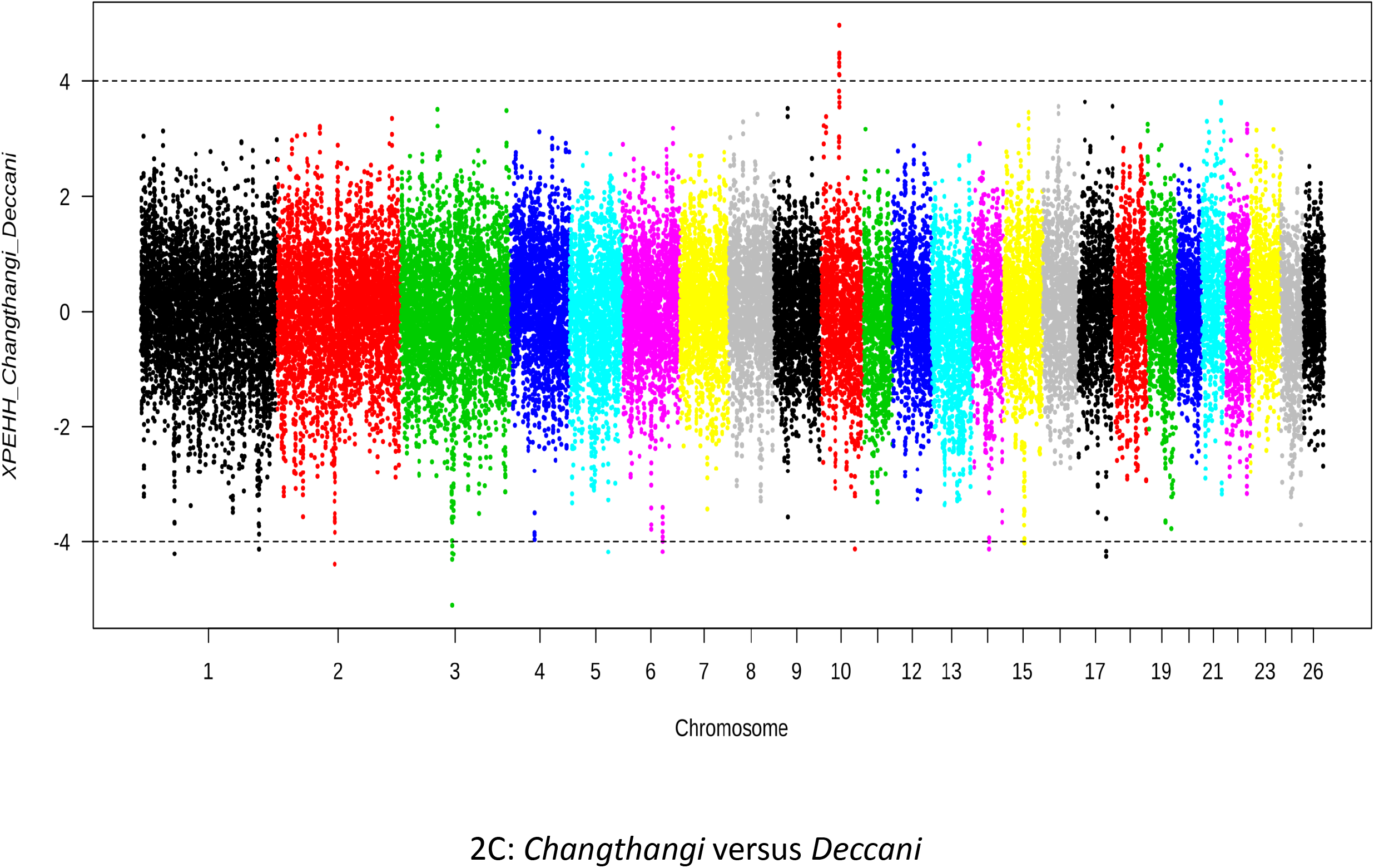
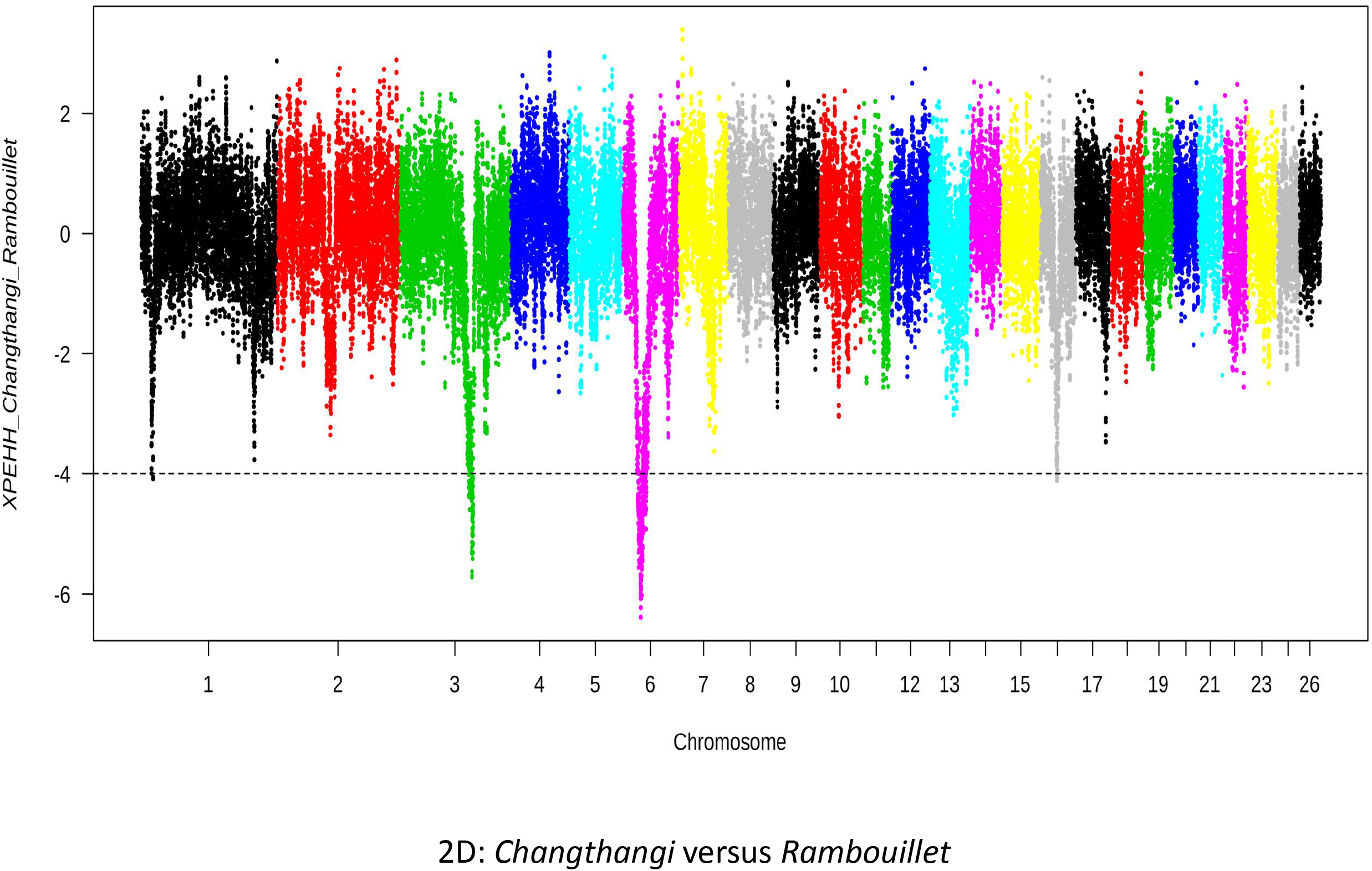
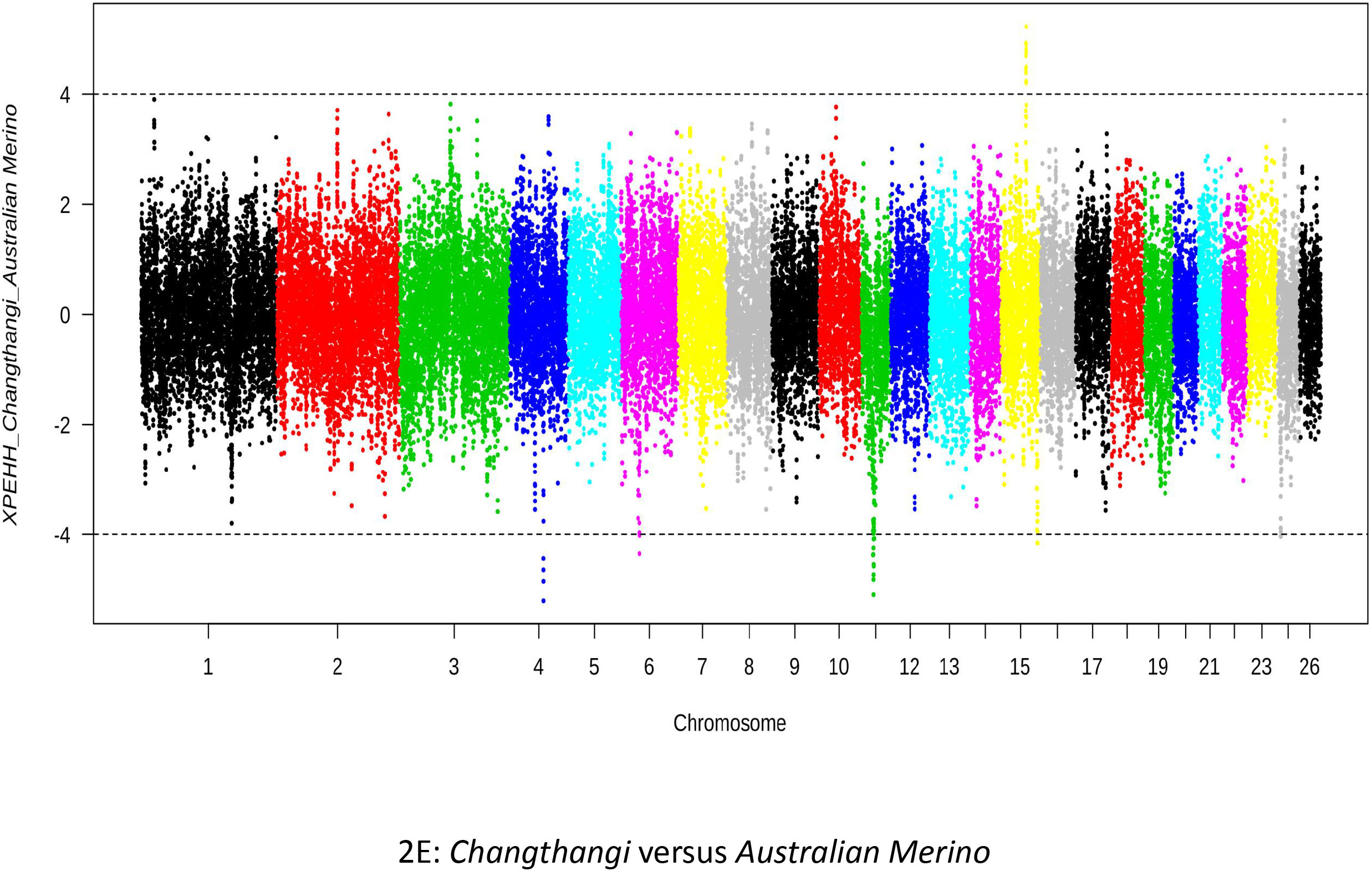
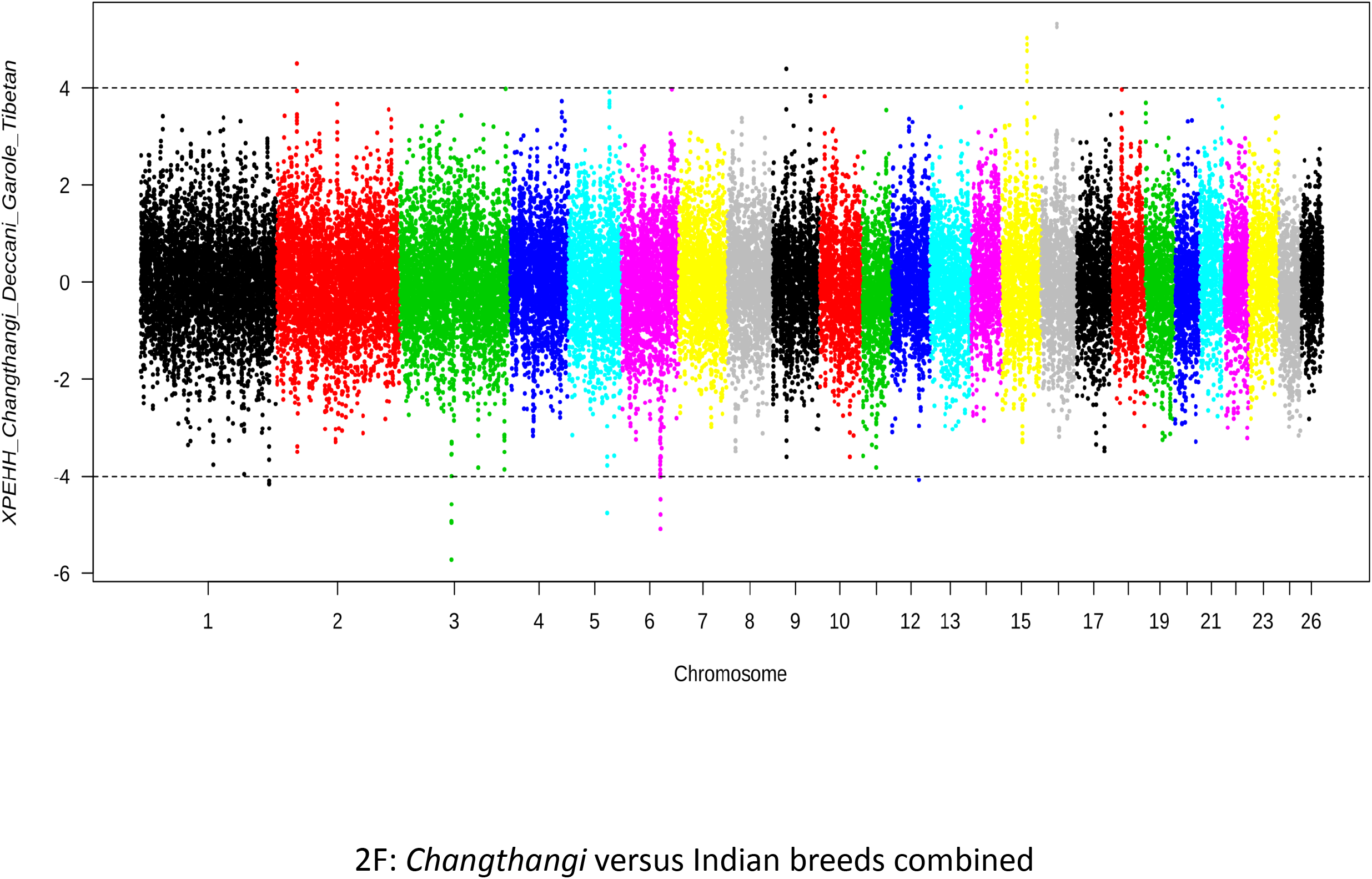
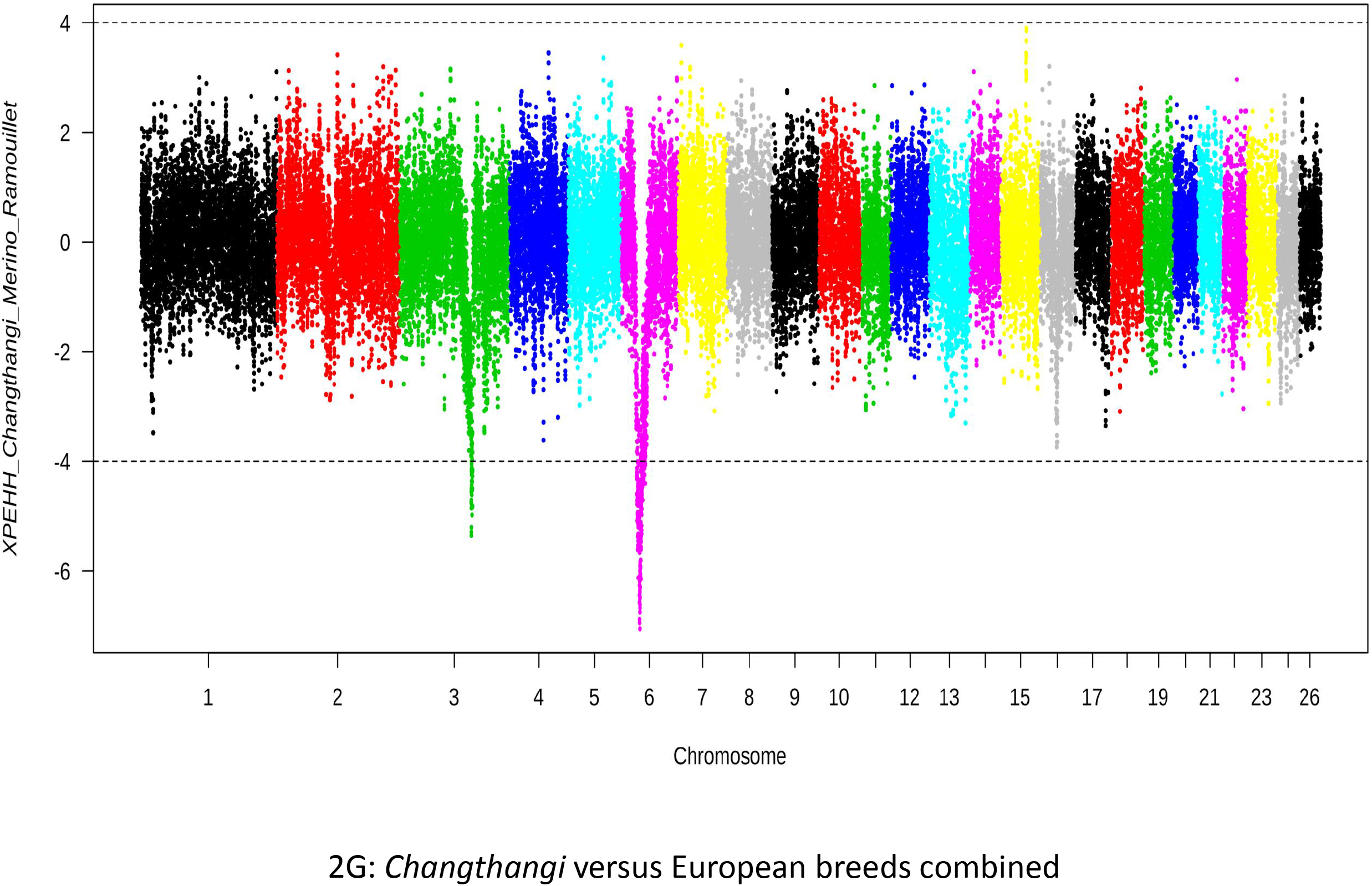
XP-EHH plots showing SNPs under selection sweeps in *Changthangi* sheep on comparison with other breeds in seven combinations.

Functional profiling using g:profiler showed the enrichment of pathways including ‘Energy Metabolism’, ‘Electron Transport Chain (OXPHOS system in mitochondria)’, ‘Exercise-induced Circadian Regulation’ and ‘Ketogenesis and Ketolysis’. Further literature mining showed that these genes are involved in stress adaptability and help in maintaining body homeostasis during extreme climatic and other environmental conditions. The UCP genes are reported to be involved in eliciting cold-induced non-shivering thermogenesis (Caron et al., 2017) which is an important adaptive character in *Changthangi* being reared and adapted to high altitude-cold climate environment. *UCP2* and *UCP3* gene expression in beige adipocytes has been reported to play an important role in cold adaptation (Pohl et al., 2019), especially in skeletal muscles in sheep (Henry et al., 2011) and beige adipocytes in pigs (Lin et al., 2017) and body metabolism (Schrauwen and Hesselink, 2002). Earlier, *UCP3* gene has also been successfully associated with growth and feed efficiency in chicken (Jin et al., 2018). *PAAF1, C2CD3* and *DNAJB13* have been reported to play important role in overall functioning of nervous system (Gao et al., 2020). Nervous system plays an important role in acclimatization of animals to cold and hypoxic conditions as perception to surrounding micro- and macro-environment and subsequent homeostatic responses are dependent on it (Tan and Knight, 2018). Heat shock proteins are also reported to play significant role in “cross-adaptation” of animals’ *vis-à-vis* cold temperatures and hypoxia (Gibson et al., 2017; Salgado et al., 2014). *DNAJB13* belongs to *DnaJ HSP40* family of heat shock proteins and is involved in body adaptation to temperature variations/ stressors (Ohtsuka and Hata, 2000). *RELT* and *PPME1* genes have earlier been reported to be related to immunoglobulin formation and thus confer resistance to pathogens especially gastrointestinal nematodes in sheep (Atlija, 2016). Similar to our results, *P2RY6* and *RELT* genes were identified in selection sweeps as candidate gene related to traits of economic importance and cold acclimation in different cattle breeds of Sweden by Ghoreishifar et al. (2020). *MRPL48* gene is related to mitochondrial function in cells which is reported to be related with thermogenesis and cold tolerance traits in different species (Cannon and Nedergaard, 2008; Li et al., 2019). Cold acclimatization is reported to be related to mitochondrial density and function in several studies (Bal et al., 2017; Dos Santos et al., 2013; Lucassen et al., 2003) and increase in the above is related to non-shivering thermogenesis in adipose tissues of animals (Hood et al., 2018; Sokolova, 2018). Similarly, *OSGEP* gene (OAR7; 23.83 Mb) has earlier been reported to be associated with mitochondrial function (Oberto et al., 2009). Based on available scientific literature, *RAB6A* gene has specific function in hypoxia-induced cellular switches and altered phenotypes (Wang et al., 2019).

Other genes under selection in *Changthangi* included *PARP2* (OAR7; 23.90 – 23.92 Mb) which was identified under selection in comparison with *Tibetan* sheep. It has earlier been found under selection in high-altitude adapted sheep breeds (*Tibetan*, *Menz* sheep and *Blackheaded Somali* population) (Edea et al., 2019; Wei et al., 2016) and associated with adaptation to hypoxic conditions (Mishra et al., 2003) in newborn piglets at high altitudes. In concurrence to the present results, *TEP1* gene (OAR7; 23.87 Mb – 23.90 Mb) found in the same comparison was identified in selection sweeps in *Blackhead Somali* and *Menz* sheep/ populations (Edea et al., 2019). We also found a region on chromosome 16 (31.70 – 31.85 Mb) containing *GHR* and *CCDC152* genes, both of which has been reported to be associated with reproductive mechanisms in sheep (Xing et al., 2019; Xu et al., 2018). Our results show the need for an in-depth exploration underlying genes and pathways involved in adaptive traits of *Changthangi* population vis-à-vis nervous system, environmental tolerance, productive and reproductive traits.

Although we were primarily concerned with the regions under selection in *Changthangi*, we also investigated the remarkable signal centred on chromosome 6 (OAR6_42834740.1; XP-EHH = −7.05) in the combined comparison with European commercial breeds (Fig). An adjacent SNP (OAR6_41936490.1, XP-EHH = −6.94) was reported to be significantly associated with body weight in *Australian Merino* sheep in a GWA study (Al-Mamun et al., 2015). This signal helps to visualize the effect of directed artificial selection on commercial breeds and provides a sharp contrast to *Changthangi*, where no such selection is practised.

To conclude, we undertook the first study to elucidate the selection sweeps of the high altitude Indian sheep breed *Changthangi*, by contrasting it with other Indian and European breeds. We found multiple functional regions on OAR 7, 15 and 16 under selection in *Changthangi* sheep which are related to adaptation to climatic stressors, hypoxic conditions and reproduction. The genes present in these regions are attractive candidates to study the functional mechanisms of adaptation to cold climate. Further, it would also be interesting to see if the regions detected in our study are under selection in other high-altitude sheep breeds of the world.

## Acknowledgements

The authors would like to acknowledge the help provided by staff of AGB division of SKUAST-K for help, support and inspiration during the study.

## Supplementary data information

**File 1:** Protein-protein interaction network of the genes under positive selection in *Changthangi*.

**File 2:** SNPs, regions and genes under selection in comparison of *Changthangi* with *Tibetan* breed.

**File 3:** SNPs, regions and genes under selection in comparison of *Changthangi* with *Indian Garole* breed.

**File 4:** SNPs, regions and genes under selection in comparison of *Changthangi* with *Deccani* breed.

**File 5:** SNPs, regions and genes under selection in comparison of *Changthangi* with *Rambouillet* breed.

**File 6:** SNPs, regions and genes under selection in comparison of Changthangi with Australian Merino breed.

**File 7:** SNPs, regions and genes under selection in comparison of Changthangi with Indian breeds combined.

**File 8:** SNPs, regions and genes under selection in comparison of Changthangi with European breeds combined.

